# Altered Glycolysis–TCA Anaplerosis Axis Impairs Monocyte Migration and Trains Macrophage Polarization in Long-term Treated HIV Infection

**DOI:** 10.1101/2025.01.15.633214

**Authors:** Alejandra Escós, Anoop T Ambikan, Sabrina Schulster, Sara Svensson Akusjärvi, Marco Gelpi, Siavash Mansouri, Prajakta Naval, Ronaldo Lira Júnior, Vikas Sood, Andreas D. Knudsen, Pratik P. Pathade, Flora Mikaeloff, Julie Høgh, Magda Lourda, Jan Vesterbacka, Piotr Nowak, Thomas Benfield, Vinay Pawar, Siegfried Weiss, Jagadeeswara Rao Muvva, Ákos Végvári, Soham Gupta, Rajkumar Savai, Susanne D. Nielsen, Ujjwal Neogi

## Abstract

Cells of the myeloid lineage, particularly monocytes and macrophages, are central to HIV pathogenesis, contributing to viral persistence and immune regulation during suppressive therapy. We hypothesized that metabolic reprogramming and altered chemokine signaling in people with HIV (PWH) on long-term ART impair monocyte trafficking and macrophage polarization. Using single-cell RNA sequencing, immunophenotyping, and metabolic modeling, we identified altered receptor expression and disrupted metabolic flux linked to reduced monocyte migration. Plasma secretome profiling revealed a nonclassical inflammatory microenvironment, while integrative multi-omics and single-cell proteomics of monocyte-derived macrophages (MDMs) demonstrated metabolic rewiring of the Glycolysis–TCA Anaplerosis Axis, orchestrated in part by elevated α-ketoglutarate (AKG). Differentiation with PWH serum or AKG, skewed MDMs toward an M2-like phenotype, and enhanced HIV susceptibility. Together, these systems-level and mechanistic analyses reveal that metabolic training drives macrophage dysfunction in well-treated PWH, sustaining low-grade inflammation and highlighting potential therapeutic targets.

## Introduction

HIV infects cells of the myeloid lineage, which play an essential role by contributing to viral replication, immune response, and maintaining immune homeostasis during suppressive HIV infection (1). Monocytes and monocyte-derived macrophages (MDM) are essential components of the myeloid lineage and play a critical role in the immune response to infection and inflammation. They can be metabolically reprogrammed to adapt to environmental changes and modulate their immune response (2). We have recently found that myeloid cells may modulate the inflammatory milieu during suppressive HIV infection due to an altered metabolic environment after prolonged successful therapy (3). However, no functional tests have been performed to confirm the direct contribution of immune metabolism to macrophage function (4). We hypothesized that metabolic reprogramming and altered chemokine signaling in people living with HIV (PWH) under prolonged antiretroviral therapy (ART) affect monocyte transport and polarization due to sustained inflammation.

The present study aimed to perform an in-depth characterization of monocyte/macrophage metabolic reprogramming to identify the mechanism of impaired function in PWH on long-term suppressive therapy. In this study, we used deep-immune profiling of metabolic transporters and chemokine receptors and single-cell RNA sequencing (scRNAseq) on circulating monocytes. We identified immunometabolic impairments of monocytes in PWH on prolonged suppressive ART compared to the people without HIV (PWoH). We examined the immunometabolic status of blood-derived MDMs by using single-cell quantitative proteomics (SCqP), advanced integrative transcriptomics, quantitative proteomics, and network analyses related to macrophage function. We characterized the metabolic rewiring by developing the macrophage-specific genome-scale metabolic model (GEM) to identify the altered metabolic flux. In addition, we investigated the functional changes in mitochondrial respiration and immunometabolism due to the microenvironment in MDMs derived from healthy blood donor monocytes with pooled serum from PWH compared to treating people without HIV (PWoH) with serum. Finally, we validated the effect of exogenous metabolite enrichment or deprivation on metabolic rewiring that caused macrophage functional impairment. Our study is the most comprehensive immunometabolic and functional characterization of MDM from well-treated PWH that identified impaired monocyte and macrophage function due to metabolic reprogramming leading to a low-grade non-classical inflammatory environment, macrophage exhaustion in long-term treated PWH and increased susceptibility for HIV-1 reinfection.

## Results

### System-Level Secretome Alterations Reveal Impaired Monocyte Migration and Immune Signaling in PWH

Previously we have shown that persistent metabolic reprogramming and altered chemokine signaling in PWH on successful therapy may impair monocyte trafficking and polarization, with downstream effects on adaptive immunity (3). Therefore, we examined plasma secretome profiles by Olink™ Explorer 3072 (Olink Ab, Sweden), and monocyte migration to assess defects in immune cell recruitment and signaling pathways between PWH and PWoH. The clinical data is presented in Table S1. Differential protein abundance (DPA) identified 489 proteins that were significantly different (adjusted p<0.05) (Table S2), of which 85% (417/489) were significantly low. The data indicate that PWH may have defective immune cell recruitment (lower levels of CCL15, CCL21, CXCL14, and CXCL17) at the inflammatory site and altered cell death process (CASP3, CASP7, CASP8, and CASP9), even though CXCL9, CCL27, and CASP2 were upregulated (Fig 1a). The cell type-specific expression profiles obtained from the human protein atlas (5) showed that most differently regulated proteins were highly expressed in the CM, IM, and NCM (Fig. 1b). Further, expression directionality incorporated gene set enrichment analysis (GSEA) identifies upregulation of hematopoietic cell lineages and cytokine-cytokine receptor interactions, indicating activation of the immune response due to the ongoing inflammatory response. In contrast, downregulation of signaling pathways such as RIG -I-like receptor signaling and FoxO signaling potentiates weak innate and adaptive immune responses against viral infections and macrophage function, supported by downregulation of Fc gamma R−mediated phagocytosis (Fig. 1c and Table S3). Earlier we also reported a significant increase in CCR5-positive cells was detected in both CM and NCM in PWH compared with PWoH (3). Increased CCR5 expression can indirectly influence monocyte migration by altering cellular behavior, signaling pathways, and the immune microenvironment. To investigate this, we isolated monocytes from PWoH (n=5) and PWH (n=19) and performed a trafficking assay in a Transwell system to measure migration in response to CCL2 (as there was no change in CCR2 expression in the monocytes between PWoH and PWH). A significant monocyte migration decreases in PWH compared with PWoH were observed (Fig. 1d). Together, these findings suggest that persistent systemic non-classical inflammatory plasma microenvironment in PWH impair monocyte trafficking and potentially impact the polarization.

**Figure 1.**
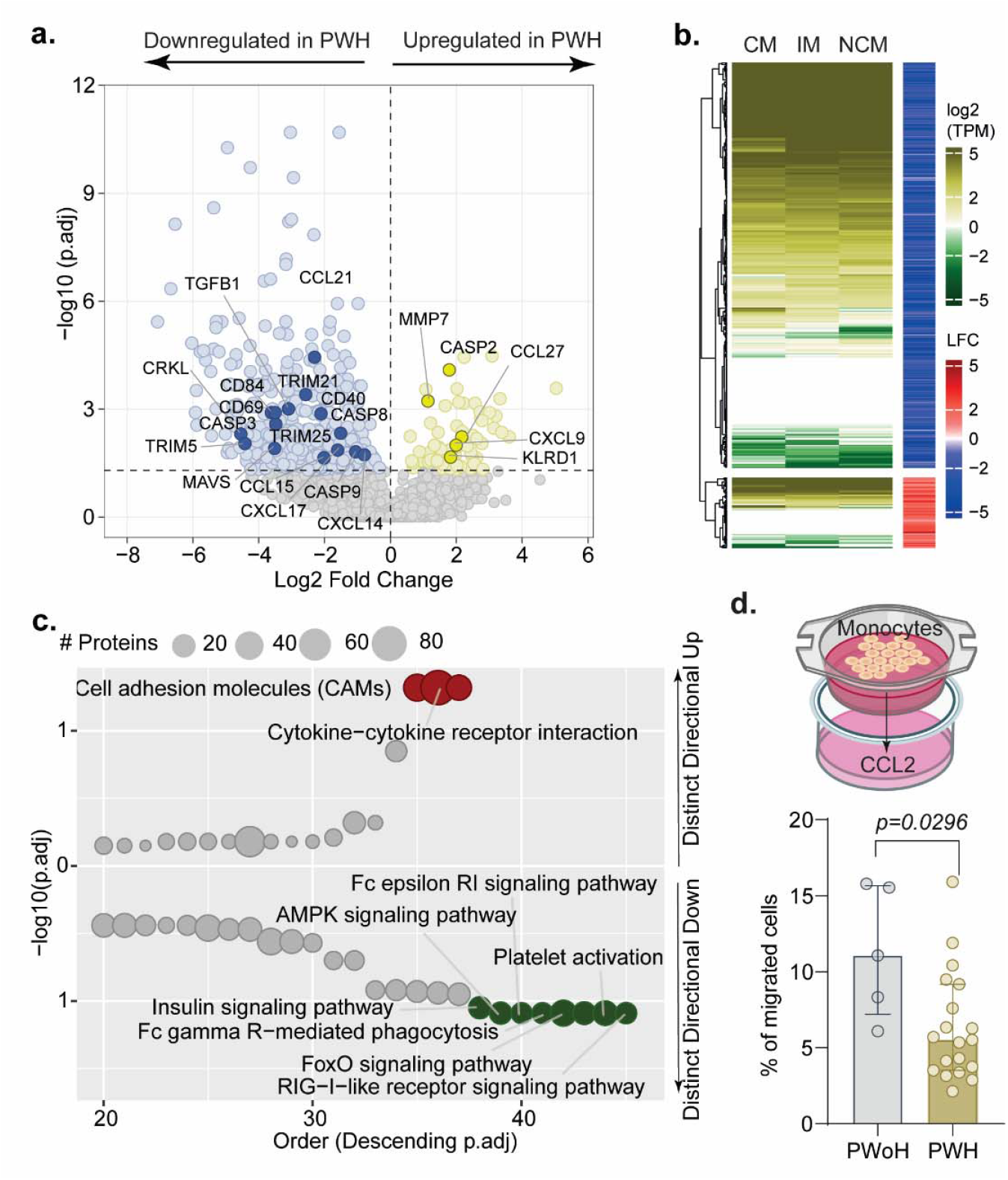
Deep plasma secretome and monocyte migration assay: **a)** Volcano diagram of differentially abundant plasma proteins between PWH and PWoH using Olink proteomics data. Proteins with adjusted p<0.05 are considered as significantly expressed in PWH compared to PWoH. **b)** The heatmap shows the expression pattern of proteins significantly regulated between PWH and PWoH in CM, IM, and NCM, which were obtained from the human protein atlas. Row annotation shows expression direction as log2 fold change according to Olink proteomics data **c)** Results of pathway enrichment analysis between PWH and PWoH using Olink proteomics data. Y-axis represents the negative log10 scaled fitted p-values belonging to a specific directional class. **d)** Migration assay scheme and percentage of migrated monocytes in the study population.

### Distinct monocyte populations and metabolic dysregulation in PWH revealed by scRNAseq

To better understand the monocyte population, we performed scRNAseq on negatively selected monocytes in PWoH (962 cells) and PWH (2856 cells). Six distinct populations of monocytes were stratified by applying the Louvain algorithm (Fig. 2a and Table S4). The populations were finalized based on the marker gene expression level and the percentage of cells in each population where they were expressed. Expression patterns of the top 15 marker genes that defined the clusters (C1 to C6) are shown in Fig. 2b. The marker genes showed unique expression profiles in the populations, confirming their distinct identity. To further study the function of each cluster, we performed GSEA of the marker genes (adjusted p<0.05 and avg log2FC>0.25) of each cluster in PWoH and PWH. Increased expression of genes involved in the PI3K-Akt signaling pathway (FOXO3, VEGFA, CDKN1A, THBS1) was observed in the PWH (C1) cluster. It is found to be absent in the PWoH (C1) cluster (Fig. 2c). Moreover, the activation of NK cell-mediated cytotoxicity was also observed in the PWH (C4) cluster, suggesting a viral suppressive nature. Analysis of the PWH (C5) cluster showed increased expression of genes belonging to pathways such as cytosolic DNA sensing, RIG-I-like receptor signaling, and NOD-like receptor signalling showing a probable presence of foreign genetic material in the cells. Further, the analysis identified that the PWH (C6) cluster is metabolically active; specifically, it showed activation of energy, carbohydrate, and amino acid metabolism pathways. Additionally, key genes in the NOD-like signalling and RIG-I-like receptor signalling pathway showed distinctive expression in the PWH (C5) cluster (Fig. 2d) in comparison with other clusters. The C5 population has increased IRF7, STAT1, and STAT2 transcription factors. IRF7 enhances IFNβ transcription during the later stages of infection, creating a positive feedback loop in which type I and type III IFNs further upregulate IRF7 expression (6). We then performed differential expression analysis to find genes regulated in each cluster of PWH compared to the corresponding clusters of PWoH. The analysis identified a total of 33 genes significantly regulated (adjusted p<0.05 and avg_log2FC>0.25) among the clusters. Of the 33 genes that differed significantly between PWH and PWoH monocyte clusters, most belonged to C2 (70%, 23/33) and were part of metabolic dysregulation (e.g., PIK3R1, LGALS2), macrophage activation (e.g., SYK), and phagocytosis (e.g., SYK and CD302) (Table S5). To identify the cluster-specific cell fate and predict the pathway of functional differentiation of monocyte clusters, we performed trajectory inference analysis by using the C1 cluster as the root. We identified a different trajectory of monocyte cell fate in PWH and PWoH (Fig. 2e). The analysis inferred trajectories of cell fate from M1 to other clusters in PWoH. In contrast, there was branching from C1 to C2 and C4 through C5 in PWH. Interestingly, the M5 cluster, in which most of the genes involved were part of the cytosolic DNA sensing pathway, was unique in PWH, indicating the presence of DNA (7) could be due to chronic immune activation. Overall, the data suggest unique cell differentiation and dysfunction in PWH, possibly due to the metabolic changes and inflammatory microenvironment resulting from low-grade persistent HIV replication. Furthermore, projection of the mean expression of genes of antigen processing and presentation on the predicted pseudo time axis of inferred trajectory starts from C1 and ends at C3 showed a dysregulated path in PWH. The path showed an inhibition from C1 to C5, then activation towards C2, followed by a drop when the path reached C3. A similar trend was also shown by the HIF-1 signalling pathway, except that the pathway showed activation from starting cluster (M1) itself (Fig 2f). On the other hand, the trajectory starts from C1 and ends at C4, showing an initial inhibition of the cytosolic DNA sensing pathway from C1 to C5 and then an activation towards C4. These findings suggest altered cell differentiation trajectories, metabolic reprogramming, and immune dysfunction in PWH, which may play a critical role in the pathogenesis of HIV-associated immune dysregulation.

**Figure 2.**
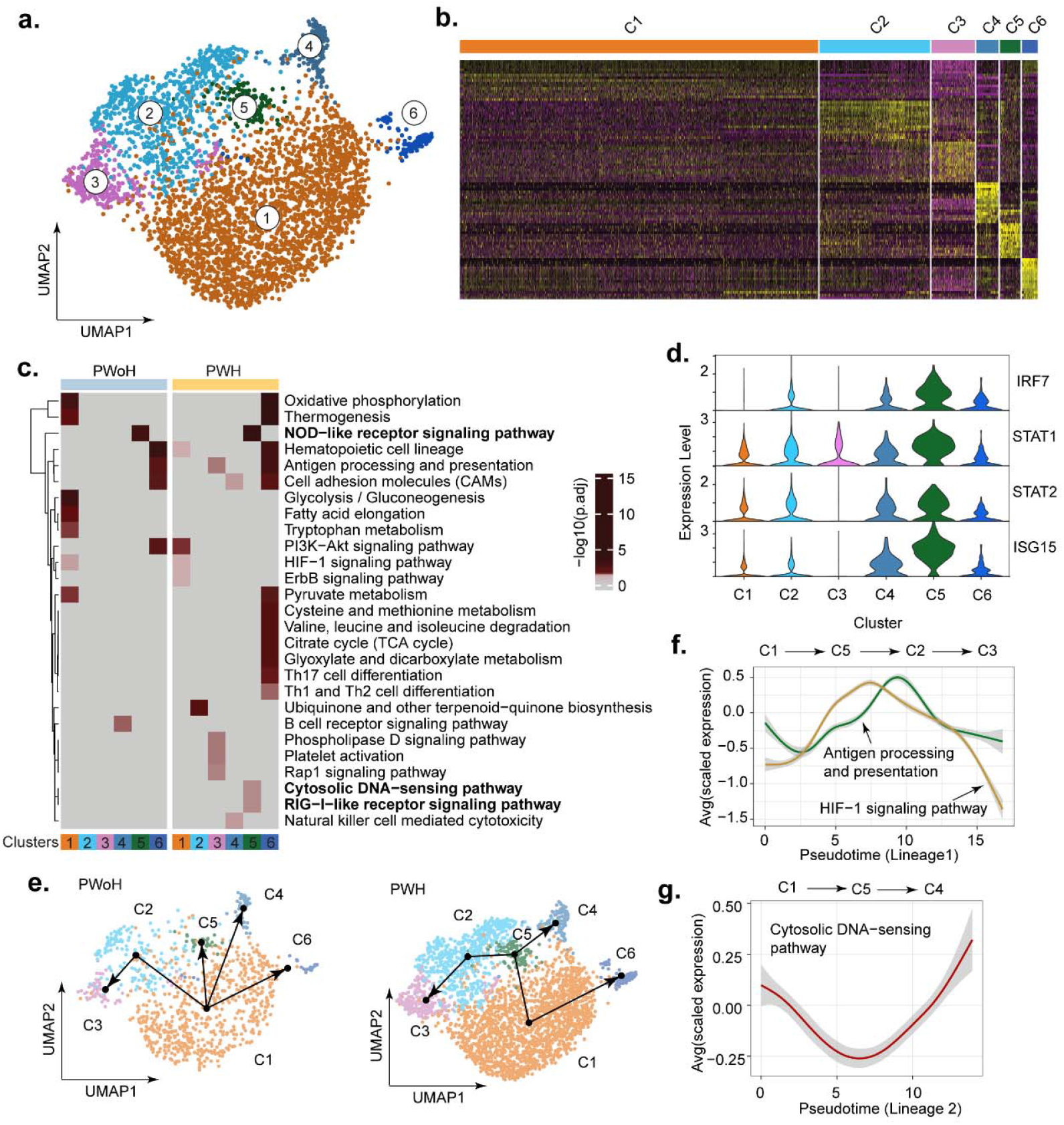
Analysis of single-cell RNA sequencing (scRNAseq) in monocytes. **a)** UMAP plot showing monocyte sub-populations identified in PWH and PWoH cells. **b)** Heatmap visualizing the expression patterns of the 15 major marker genes of each monocyte cluster. The row labels represent the monocyte clusters. **c)** Significantly enriched signaling pathways in each monocyte cluster. Heatmap shows negative log10-scaled adjusted p-values of pathway enrichment. Column labels show monocyte clusters. The color of the text indicates unique pathways for each cluster colored according to the cluster. d) Violin plot visualizing the expression of selected significant genes from NOD-like and RIG-I-like receptor pathways in the monocyte sub-populations in PWH. **e)** Trajectory of functional differentiation of monocytes derived from single cell transcriptomics data. The line connecting the clusters represents the derived trajectory. **f)** Dynamics of average expression of genes in antigen processing and presentation and HIF-1 signalling pathway projected onto the predicted pseudotime of first lineage (C1-C5-C2-C3) in PWH. **g)** Dynamics of average expression of genes in cytosolic DNA sensing pathway projected onto the predicted pseudo time of second lineage (C1-C5-C4) in PWH

### Single-cell proteomics reveal hyperactive immunometabolism and homogeneous macrophage phenotypes in PWH

The systemic secretome (Fig 1) and scRNAseq (Fig. 2) revealed potential monocyte immune activation and an inflammatory microenvironment in PWH. Therefore, we differentiated monocytes into MDM for seven days using the granulocyte-macrophage colony-stimulating factor (GM-CSF) to understand the macrophage phenotype better (Fig 3a). We performed liquid chromatography-mass spectrometry (LC-MS /MS)-based quantitative single-cell proteomics (SCqP) on 288 MDMs from eight PWHs and compared them with 144 MDMs from four PWoH to understand the cellular heterogeneity of macrophages at the single-cell level. We quantified 993 proteins. The upregulation of metabolic genes such as isocitrate dehydrogenase 1 (IDH1), malate dehydrogenase 1 (MDH1), arginase 1 (ARG1), and glutamate dehydrogenase 1 (GLUD1) identified by DPA indicate increased metabolic activity and changes in energy metabolism. Downregulation of cytochrome c oxidase (COX), acetyl-CoA acetyltransferase 2 (ACAT2), superoxide dismutase 2 (SOD2), interferon-gamma-inducible protein 30 (IFI30), and human leukocyte antigen (HLA) indicates alterations in lipid metabolism, decreased antioxidant defense, or impaired immune cell function (Fig. 3b and Table S6). Gene set enrichment analysis further supports this (Fig. 3c). Next, we performed K-Means-based clustering to determine the similarity of cells with respect to the protein expression profile and function and identified three cell clusters (Fig. 3d). Interestingly, the MDMs of PWH were more homogeneous than the MDMs of PWoH (Fig. 3e), with the cells of cluster 2 showing the greatest homogeneity (Fig. 3e). Homogeneity of the cells were quantitatively accessed by computing the distance from centroids for each cell in each cluster. The assessment showed that cluster 2 is more homogeneous compared to others (Fig 3f) and their functions were mainly related to metabolic processes (Fig. 3g). Based on this data we hypothesized that the hyperactive immunometabolic states of monocytes lead to a depleted MDM phenotype with impaired polarization.

**Figure 3.**
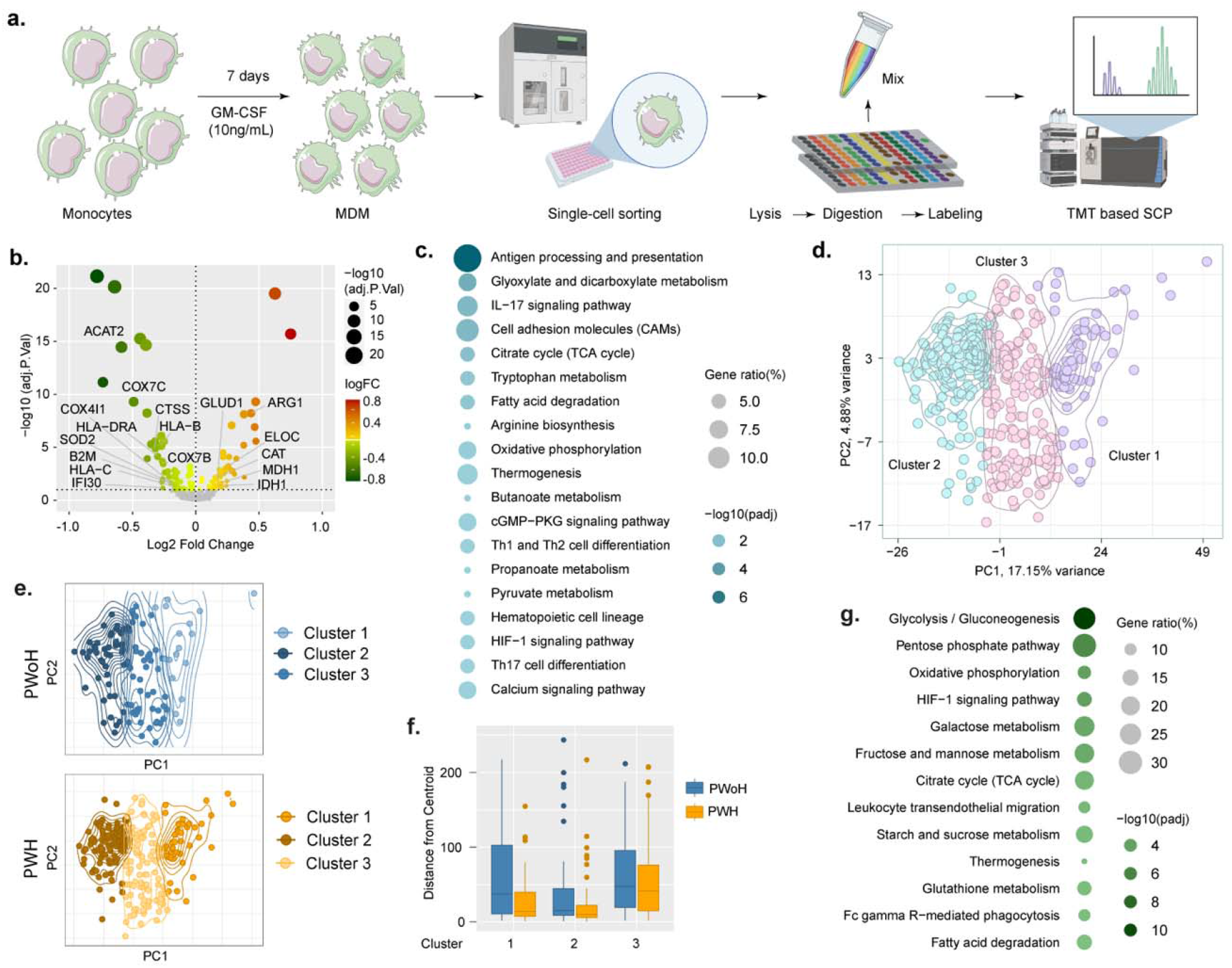
Analysis of single-cell quantitative proteomics (SCqP) in MDM. **a)** Experimental scheme of differentiation of blood-derived monocytes to MDM. **b)** Volcano diagram of differentially abundant proteins between PWH and PWoH using single-cell proteomics data. Bubble size is relative to the negative log10-scaled fitted p-values, and the color gradient corresponds to the log2-scaled fold change in expression level. **c)** Results of pathway enrichment analysis of PWH vs PWoH using single cell proteomics data. The bubble plot shows significantly enriched pathways (adjusted p < 0.1) in PWH. Bubble size is relative to the ratio of proteins significantly regulated in the corresponding pathway. The color gradient is relative to the negative log10-scaled adjusted p values of pathway enrichment. **d)** Cell clusters were identified by applying the K-Means algorithm to single-cell proteomics data. The distribution of cells is visualized in PCA coordinates. **e)** Visualization of cell distribution of the three K-means clusters identified in PWH and PWoH. The distribution of cells is visualized in PCA coordinates. **f)** Boxplot showing the variation of homogeneity between cells in PWH and PWoH. The Y axis indicates the distance of each cell from the centroid of the cluster. **g)** Results of pathway enrichment analysis using proteins uniquely expressed in cell cluster 2. The bubble plot shows significantly enriched pathways (adjusted p<0.05). Bubble size is relative to the ratio of unique proteins in cluster 2 in the corresponding pathway. The color gradient is relative to the negative log10-scaled adjusted p values of pathway enrichment.

### Metabolic modeling of MDM shows altered Glycolysis–TCA anaplerosis axis orchestrated by AKG

We developed MDM-specific GEM by integrating the MDM RNAseq data and the supernatant metabolomics based on our recently described approaches (7). We identified 253 unique reactions in PWH MDMs that revealed significant flux changes in the transport of amino acids and TCA cycle intermediates (Fig. 4a and Table S7). The unique reactions that have opposite predicted flux are presented in Fig 4b, indicating an activated and broken TCA cycle and AKG production from oxalosuccinate (Fig 4c). Specific metabolite ratios that modulate macrophage polarization are AKG/Isocitrate and Fumarate/Succinate ratios. AKG/Isocitrate ratio is key to generating itaconate, an important anti-bacterial agent that was not significantly altered (Fig 4d). On the other hand, PWHs had a decreased Fumarate/Succinate ratio, which is linked to increased OXPHOS (Fig 4d). The ratio of glutamate/glutamine was found to increase significantly (Fig 4d) in PWH, which has been described as impeded by anti-HIV-1 CD8+ T cell response (8). To universalize our findings, we performed a meta-analysis of the plasma metabolomics with 375 patients from four different countries that were reported by us previously (3, 9, 10) by comparing the PWoH (n=100) and PWH on ART (n=275). There were 216 metabolites were significantly differently abundant (adj p<0.05). We found that both AKG and glutamate were significantly elevated in PWH compared to the PWoH apart from TCA-cycle intermediate aconitate, aspartate and fumarate and glycolysis product pyruvate and lactate while decrease in citrate (Fig. 4e, Table S8), indicating that this is a universal phenomenon in PWH on therapy. We therefore hypothesize that glutamate/AKG-driven microenvironment impairs the macrophage function by altering the Glycolysis–TCA anaplerosis axis. To identify the role of the plasma microenvironment, first, we differentiated donor monocytes with either pooled serum from PWH on ART or PWoH (11) and measured the oxygen consumption rate (OCR) with the XF-Seahorse extracellular flux analyzer. The data showed an increase in OCR in MDM differentiated with the pooled serum from PWH than PWoH in all states of mitochondrial capacity (Fig. 4f). At the same time, intracellular levels of glutamate were found to have an increase trend in MDM due to the microenvironment of PWH serum (Fig 4g). These results demonstrate that dysregulation of the MDM metabolic profile in PWH could also be achieved by the microenvironment. We also measured the glutathione oxidation ratio and found that the predicted increase of oxidation of glutathione was also evident in our laboratory assay (Fig 4h). PWH had lower levels of reduced glutathione (GHS) than PWoH (Fig 4h). This indicates that the PWH plasma microenvironment activates the glutathione metabolism by increasing the expression and activity of critical enzymes involved in the glutathione redox system. To mimic the effect of increased AKG in plasma in macrophage, we differentiated MDM from PWoH with and without dimethyl-α-ketoglutarate (DM-AKG, which is permeable for AKG). We measured intracellular levels of glutamate as a key marker of immunometabolic reprogramming and observed an increase in glutamate levels (Fig 4i). We then analyzed the ratio of [GSH]/[GSSG] as another hallmark for AKG and observed that differentiation with DM-AKG increased the oxidation of GSH (Fig 4j). Overall, these data demonstrate that the altered metabolic environment in PWH on ART drives the immunometabolic reprogramming of MDMs, specifically through elevated AKG levels.

**Figure 4.**
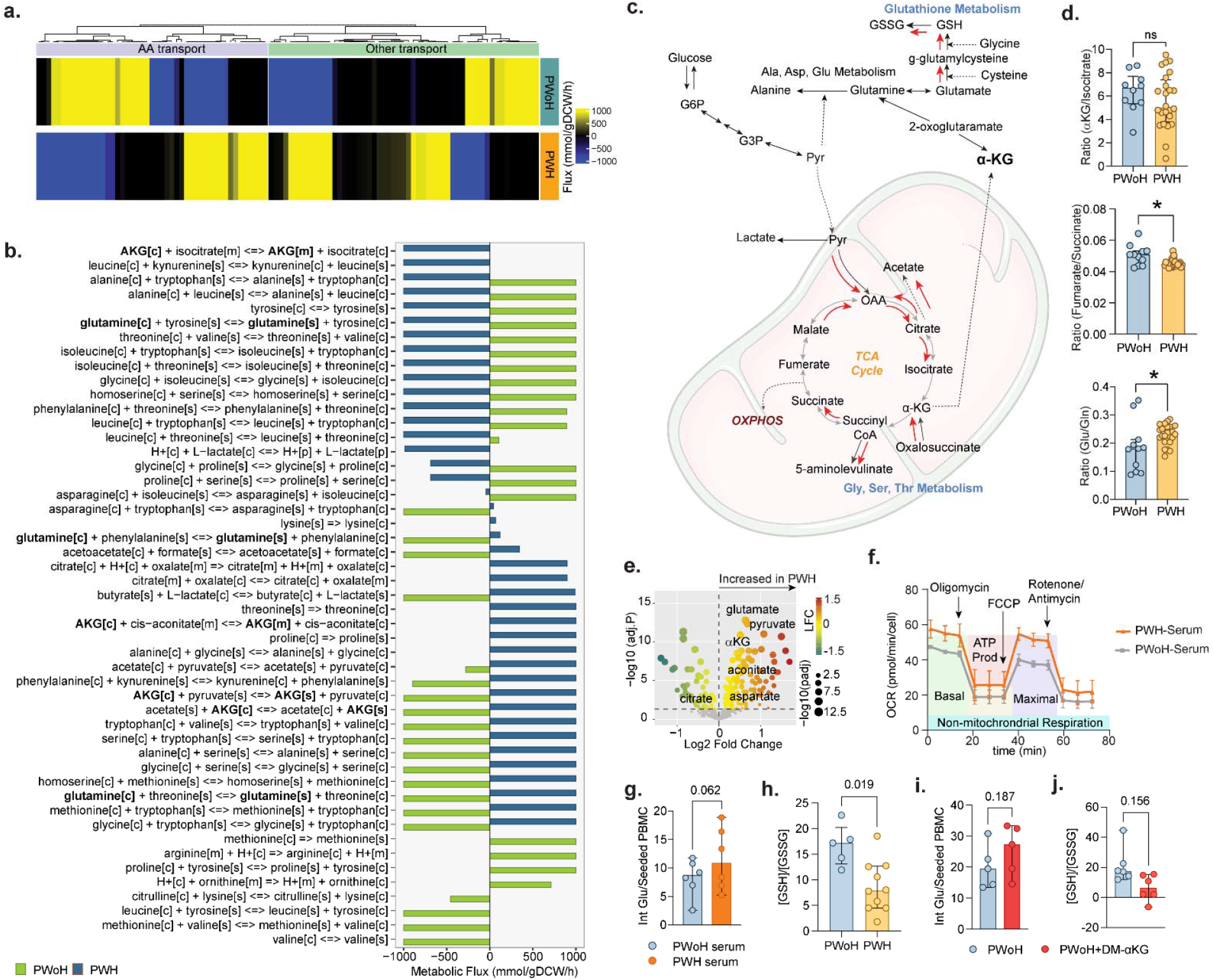
Metabolic modeling and exogenous metabolite accumulation or deprivation in MDMs. **a)** Results of flux balance analysis using MDM transcriptomics data. The heat map visualizes the predicted flux values of transport reactions in PWH and PWoH. Column labels represent the type of metabolites transported in the corresponding reactions. **b)** Bar chart showing the differences in the metabolic flux of reactions transporting amino acids and TCA cycle intermediates in PWH and PWoH. c) Reaction diagram illustrating the direction of reactions belonging to different metabolic processes. The red colored arrow indicates the direction of reactions in PWH. **d)** Metabolite levels in the plasma of PWH (n = 24) and PWoH (n = 11). **e)** Volcano diagram of differentially abundant metabolites between PWoH and PWH on ART using plasma metabolomics data. Bubble size is relative to the negative log10-scaled fitted p-values, and the color gradient corresponds to the log2-scaled fold change in expression level. **f)** Mitochondrial oxygen consumption rate (OCR) and maximal respiratory capacity of donor monocytes differentiated to macrophages (n = 5) with the pooled serum of PWH or PWoH. **g)** Intracellular glutamate in MDM differentiated (n = 6) with pooled serum from PWoH or PWH. **h)** Glutathione reductase ratio in MDM from (n = 5) PWoH or (n = 10) PWH. Statistical analysis was performed using the Mann-Whitney U test or Wilcoxon test (significance level, p<0.05) and presented with a median with 95% CI. **i)** Intracellular levels of glutamate as a key marker of immunometabolic reprogramming **j)** The ratio of [GSH]/[GSSG] in donor samples treated or untreated with DM-αKG

### Impaired macrophage function and polarization in PWH

As we identified the dysregulation of the immunometabolic homeostasis that leads to increased OXPHOS and altered glutathione metabolism, we hypothesized that the impaired immunometabolic alterations led to macrophage dysfunction. Therefore, we performed bulk transcriptomics and quantitative proteomics analysis to identify changes in macrophage polarization and function in MDM in PWH (n=24) and compared them with PWoH (n=12). The differential gene expression analysis of the transcriptomics data identified 1688 genes with significant differential regulation between PWH and PWoH (adjusted p<0.05). (Table S9). Most proinflammatory markers, e.g., TNFα, CXCL8, or STAT1, were downregulated in PWH (Fig. 5a). In the analysis of anti-inflammatory markers, we observed a significant decrease in STAT3, IL13, IRF4, or CCL4, but in contrast, some genes were significantly upregulated, e.g., PPARG and IL1R1 (Fig. 5b). The quantitative proteomics analysis identified 881 proteins significantly regulated in PWH (adjusted p<0.05), and showed a similar pattern with downregulation of most pro-inflammatory M1 and anti-inflammatory M2 markers (Fig 5c) as observed in the transcriptomics data (Fig 5a and 5b). The upregulation of anti-inflammatory M2 markers VCAM1 (recruitment and adhesion of immune cells to sites of inflammation), MMP19 (tissue remodeling processes), STAB1 (involved in tissue repair, immune regulation, and resolution of inflammation), CSF1R (essential for organogenesis), etc. (Fig. 5c and Table S10), however, suggests potentially altered recruitment, adhesion, differentiation, and proliferation of immune cells due to macrophage dysfunction. One of the functions of macrophages is the secretion of cytokines and chemokines for immunoregulation. We then measured pro-inflammatory and anti-inflammatory cytokines and chemokines secreted to the supernatant. Olink™ 96 inflammation identified a general decrease in the cytokine-secreted pro-inflammatory M1 cytokines and chemokines, e.g., TNF-a, CXCL9, and CXCL10, and anti-inflammatory M2 cytokines, e.g., IL10 and IL13 (Fig 5d). This general decrease was also observed in deep secretome analysis of plasma using the Olink™ Explorer 3072 (Olink Ab, Sweden). To explore more functions of macrophages, we assessed their phagocytosis capacity in macrophages from PWH (n = 10) and PWoH (n = 13). We observed a non-significant increase in the phagocytic capacity of MDM from PWH. To evaluate the influence of the microenvironment on macrophage function, we differentiated donor monocytes into macrophages using pooled serum from either PWH or PWoH. These macrophages were then polarized into M1 (IFN-γ and LPS) or M2 (IL4) for 24h. As expected, M1 macrophages exhibited lower phagocytosis levels compared to M2 macrophages (Fig 5f). When analyzing the phagocytosis capacity of macrophages exposed to pooled serum from PWH or PWoH, we found increased phagocytosis in both M1 and M2 macrophages, with statistical significance observed only in M1 macrophages (Fig 5g). These findings suggest that macrophage function is impaired in PWH undergoing prolonged therapy, regardless of whether the macrophages are derived directly from the individual or influenced by the microenvironment.

**Figure 5.**
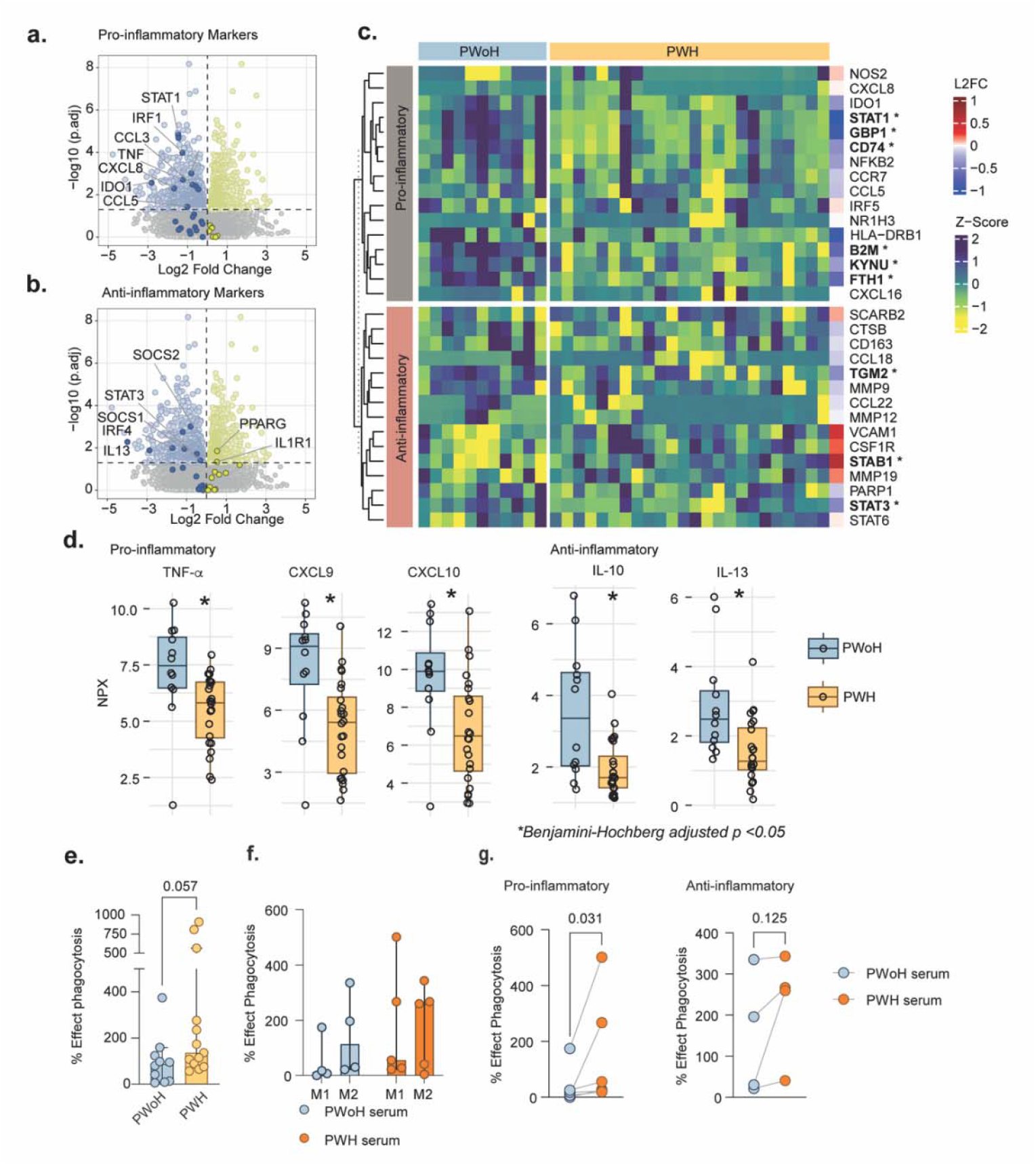
MDM proteomics analysis of macrophage polarization genes. Expression status of **a)** proinflammatory and **b)** anti-inflammatory marker genes in PWH compared with PWoH based on MDM transcriptomics data. Marker genes are labeled in the PWH vs PWoH volcano plot. **c)** Expression landscape of pro- and anti-inflammatory marker genes based on MDM proteomics data. The heatmap visualizes the z-scaled MDM proteomics expression values of the marker genes. The row labels of the heatmap show the fold change values of the marker genes in PWH compared to PWoH. **d)** Representation of Olink proteomics data pro-inflammatory and anti-inflammatory cytokines and chemokines shown as fold-change values of protein expression in PWH compared with PWoH. Significant expression (adjusted p < 0.05) is indicated with an asterisk. Phagocytosis assay of E. coli **e)** from PWH (n = 13) and PWoH (n = 10) MDM, **f)** and **g)** from donor MDM (n = 6) first differentiated with pooled serum of PWoH or PWH and afterwards towards pro-inflammatory M1 (LPS and IFN-g) or anti-inflammatory M2 (IL4). Statistical analysis was performed using the Mann-Whitney U test or Wilcoxon test (significance level, p<0.05) and presented with a median with 95% CI.

### AKG-driven metabolic reprogramming in PWH promotes M2-like macrophage polarization and increased HIV susceptibility

Classical macrophage polarization depends on metabolic changes that control macrophage function (12). Based on the above results, we hypothesized that macrophages from PWH were “trained” or polarized differently because of the metabolic reprogramming created by prolonged therapy, nonclassical inflammation, and immune activation. Our analysis of their phagocytic capacity revealed that, while not statistically significant, PWH macrophages exhibited increased phagocytosis in their non-polarized state, suggesting a tendency toward M2-like polarization. We developed a FACS panel to characterize macrophage polarization and measured levels of CCR5 and CD4, both HIV-1 entry receptors (Fig Sup 1a). We differentiated PWoH macrophages into non-polarized, M1 macrophages or M2 macrophages. Macrophages with M1-like phenotype had higher MFI levels of HLA-DR within the CD64^+^/CD86^+^ population (Fig. Sup 1b). Macrophages with M2-like phenotype were characterized by CD200R^+^ and/or CD163^+^ cells (Fig. Sup 1c). These findings confirm that the phenotype of macrophages from PWH in their unpolarized state resembles more M2-like macrophages (Fig 6a). This trend was also influenced by the microenvironment and DM-AKG differentiation, though the M2-like phenotype was less pronounced in pooled HIV serum-differentiated macrophages (Fig 6c and 6e). A decrease in CD86 MFI in PWH further supports the presence of an M2-like phenotype. However, this decrease was not observed in macrophages differentiated with pooled HIV serum or DM-AKG (Fig. Sup 1d, 1e, and 1f). This indicates that while AKG does not fully alter macrophage function, it plays a significant role in the shift towards M2-like polarization. The CCR5 receptor and CD4 co-receptor are critical for HIV infection, so we measured their expression levels on the cell surface. In this instance, we observed a significant increase of CCR5 and CD4 expression levels in macrophages from PWH, indicating these cells are more likely to be susceptible to HIV infection (Fig 6b and Fig Sup 1d). Examining the impact of the microenvironment, we found that CCR5 levels were significantly elevated in macrophages differentiated with either pooled HIV serum or six individual serums from PWH (Fig 6d and Fig Sup 1e, 1g). In contrast, CD4 co-receptor levels did not show significant changes, although there was a general trend toward an increase in macrophages differentiated with serum from PWH (Fig Sup 1e and 1g). These findings suggested an increased susceptibility for HIV-1 spread. To explore this further, we conducted a time-course analysis of HIV-1 production over 14 days in donor MDM (Fig Sup 2a). Viral production began consistently on day 5, peaked on day 7, and experienced a second wave on day 10 (Fig. Sup 2a). Both days 7 and day 14 showed high percentage of intracellular Gag^+^ macrophages (HIV-1, infected cells), but cell death increased significantly by day 14 (Fig Sup 2b). Macrophages differentiated with DM-AKG exhibited significantly higher HIV infection rates, which correlated with increased CCR5^+^ receptor expression (Fig 6f). This suggests that higher AKG levels promote increased HIV infection and spread in PWH, making HIV-1 persistence more challenging if AKG levels aren’t regulated.

**Figure 6.**
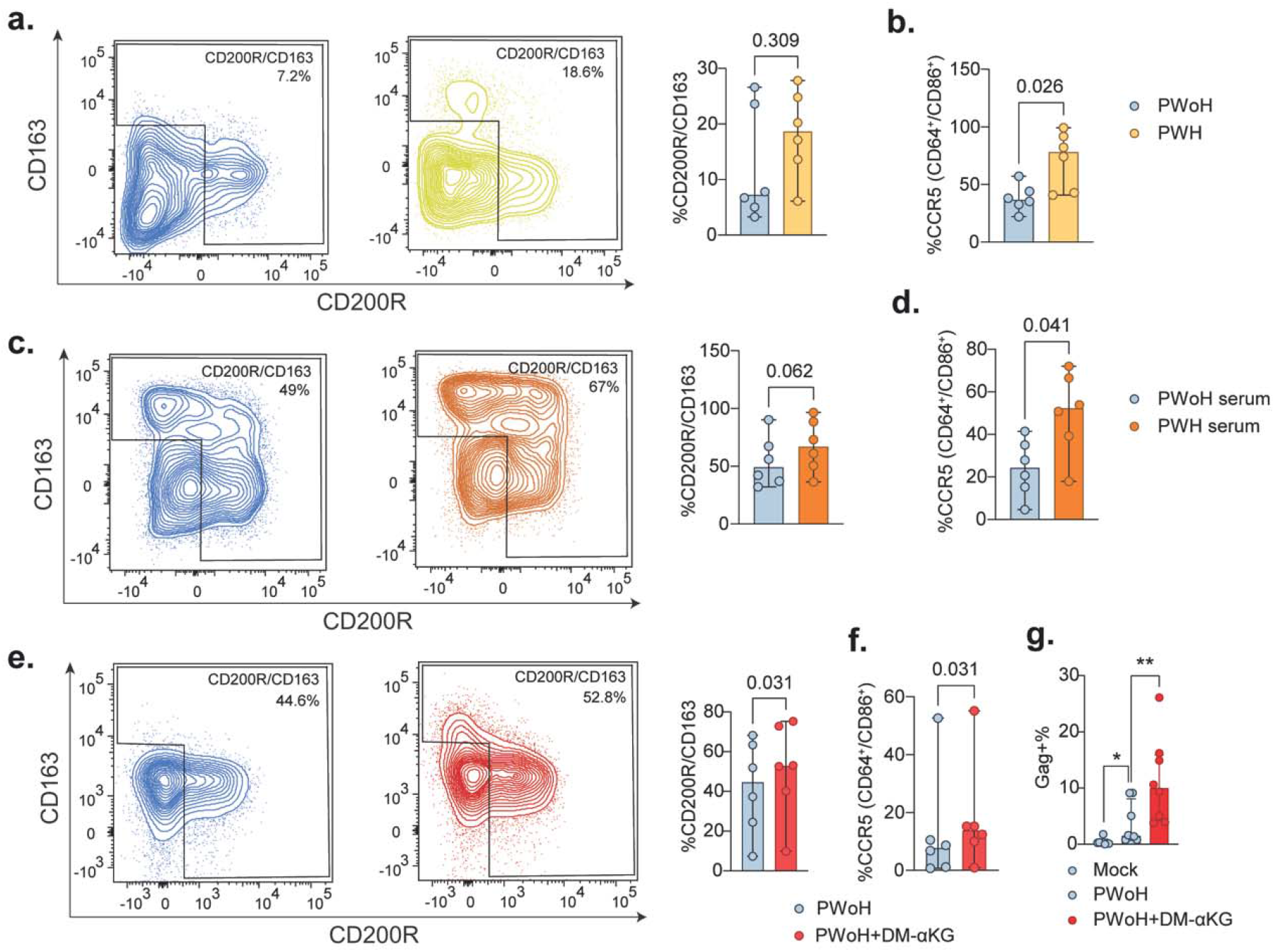
M2-like macrophage polarization and HIV infectivity. **a)** Contour plots and graphs showing the percentage of cells expressing CD200R and CD163 in PWoH (n = 6) and PWH (n = 6) and **b)** representation of CCR5 percentage. **c)** Contour plots and graphs showing the percentage of cells expressing CD200R and CD163 from donor MDM (n = 6) differentiated with pooled serum of PWoH or PWH and **d)** representation of CCR5 percentage. **e)** Contour plots and graphs showing the percentage of cells expressing CD200R and CD163 from donor MDM (n = 6) differentiated with or without DM-AKG and **f)** representation of CCR5 percentage. The contour plots show a representative sample in each group with the median expression level within that group. **g)** HIV-1_ADA-M_ Gag^+^ proportion of non-infected and infected with HIV-1_ADA-M_ of donor MDM (n = 6) differentiated with or without DM-AKG. Statistical analysis was performed using the Mann-Whitney U test or Wilcoxon test (significance level, p<0.05) and presented with a median with 95% CI.

## Discussion

Our study explored the dysfunction of monocytes and macrophages in well-treated PWH. We identified impaired monocyte migration in a subset of PWH. We found that macrophage polarization induced by metabolic reprogramming contributed to a nonclassical, para-inflammatory microenvironment along with heightened monocyte activation. Additionally, our data revealed impaired cytokine regulation *in vitro* within PWH-derived MDM, linked to macrophage exhaustion. Moreover, the depletion of anti-inflammatory macrophages may promote an apoptosis-resistant phenotype, which in turn drives inflammatory cascades in PWH. Finally, macrophage immunometabolic reprogramming was found to be driven by Glycolysis–TCA anaplerosis axis potentially due to the elevated AKG, which, in consequence, changed macrophage polarization towards M2-like and increased HIV-1 infection *ex vivo*.

Interestingly, analysis of cell fate trajectories identified a bifurcation pathway of monocyte differentiation by genes involved in DNA sensing pathways unique to PWH. Studies have reported that prevention of HIV integration by integrase inhibitors such as raltegravir leads to the formation of HIV-1 episomal DNA (2-LTR) during suppressive therapy (13), which can activate the DNA sensing pathway (14). Once they bind to the foreign DNA, the DNA sensors initiate a cascade of signals that activate the monocytes, typically involving the production of type I interferons and pro-inflammatory cytokines (15), which can influence monocyte trafficking as observed less trafficking capacity of monocytes in PWH *in vitro*. The earlier study also indicated that the transition from monocytes to macrophages is an instructive process that occurs through canonical monocyte chemokine receptor signaling and primarily regulates monocyte traffic and monocyte fate during inflammation (16). We also reported an increase in CCR5^+^ monocytes in PWH (3). *In vitro* transmigration assays showed that the CCR5 contributed to monocyte spreading in shear flow (17) and thus may influence monocyte recruitment (17, 18).

Consistent with scRNAseq and immunophenotyping data, our in-depth secretome analysis revealed higher levels of CXCL9 and CCL27, which play essential roles in immune cell recruitment and inflammation (19). However, we also observed lower levels of CD84 and CD69, which play a role in monocyte and macrophage activation and differentiation (20, 21). Further low CXCL17, CCL21, CCL15, and CXCL14 may decrease the accumulation of immune cells at inflammatory sites (22). No differences were found in the classical inflammatory molecules such as IL-6 and CXCL10. Therefore, the inflammatory state in PWH represents a state of non-classical chronic para-inflammation (23), an adaptive response of the immune system to oxidative stress leading to inflammaging (24). Interestingly, earlier studies reported that peripheral inflammation increased CCL5/CCR5 axis activation, disrupted the blood-brain barrier in mice (25), and contributed to the pathogenesis of atherosclerosis (26), which is common in PWH.

The sequel of the monocyte maturation to the monocyte-derived macrophage process is often heterogeneous (27) and microenvironment-driven. *In vitro*, studies highlighted the metabolic plasticity of the macrophages, which can be rewired in their cellular metabolism in the context of pro- or anti-inflammatory polarization and often do not fully fit in the binary framework (28). Our SCqP of MDM identified a trained MDM population in PWH with upregulation of metabolic proteins such as ARG1, IDH1, and MDH1, indicating metabolically dysregulated, hyperactive MDMs. The increased cellular metabolism identified in the transcriptomics and proteomics analysis of PWH MDMs resulted in impaired macrophage polarization. This could be due to the microenvironment, as healthy MDMs treated with PWH serum exhibited increased cellular metabolism due to increased energy demand, as the microenvironment is often observed to control macrophage function (29).

In addition, the increase in amino acids serine, valine, isoleucine, and leucine from glycolysis and the decrease in G6P in the supernatant of PWH MDM reflect a shift in glucose metabolism away from glycolysis and toward the alternative serine biosynthetic pathway and branched-chain amino acid (BCAA) metabolism. We also observed a decrease in TCA intermediates (aconitate, fumarate, and malate), reflecting a shift in cellular metabolism away from the TCA cycle and toward other metabolic pathways, such as the pentose phosphate pathway (PPP), which was supported by an increase in the G6PD transcript in PWH MDM. These data suggest defects in cellular energy metabolism associated with an impaired ability of macrophages to respond to secondary stimulation, resulting in ‘immunometabolic paralysis’ of macrophage function (2), potentially due to mitochondrial dysfunction (30). The MDM context-specific GEM further identified the activation of metabolic responses in mitochondrial fatty acid oxidation, the TCA cycle, and AKG production from oxalosuccinate that activated glutathione metabolism by increasing the expression and activity of key enzymes involved in the glutathione redox system (31). This data highlight a broad rewiring of the Glycolysis–TCA Anaplerosis Axis, where impaired glycolytic flux, reduced TCA activity (32) collectively reflect a state of metabolic inflexibility, which could lead to an exhausted macrophage phenotype.

An increase in AKG levels can lead to glutamate production (33), as observed in PWH plasma reported by us in several cohorts of treated PWH (3, 10, 34), and play an essential role in comorbid metabolic syndrome in PWH (9). A unique phenomenon of increased production of intracellular glutamate was observed in PWH MDM after AKG elevation altered the balance in cytokine signaling cascades. We observed a depleted anti-inflammatory phenotype when we stimulated differentiated MDM with IL-4 using PWH serum. M2 macrophages are particularly effective at efferocytosis and contribute to the resolution of inflammation (35). The plasma secretome data showed that the upregulation of CASP2 acts as an initiator caspase in the intrinsic (mitochondrial) pathway of apoptosis, and downregulation of CASP3, CASP7, CASP8, and CASP9 could cause dysregulation in the apoptotic pathways (36). Similar trends were observed in the transcriptomics and proteomics of MDM. While CASP2 may be more sensitive to cellular stressors, downregulation of the other caspases may impair the cell’s ability to undergo apoptosis in response to these stress signals, leading to cell survival, termed apoptosis resistance (37), which mediates the inflammatory cascades leading to chronic para-inflammation in PWH.

Our study has several limitations that merits comments. First, the scRNAseq analysis was performed on a limited number of PWoH samples, which constrains the statistical power and generalizability of the cell-state differences observed. Second, although we identified multiple metabolic perturbations, we focused on AKG as a proxy metabolite to probe macrophage reprogramming because of its central role at the intersection of the TCA cycle, amino acid metabolism, and epigenetic regulation. We acknowledge that other metabolites, including glutamate and lactate, may also contribute to the observed phenotypes and warrant further investigation. Finally, while our *in vitro* mechanistic studies cannot fully recapitulate the *in vivo* complexity of tissue macrophages, they mirror the high-throughput multi-omics data and provide functional support for the immunometabolic rewiring observed in PWH. Despite these limitations, our study has several notable strengths. First, we combined multi-omics profiling at the systemic, cellular, and single-cell levels with functional assays, allowing us to capture system-level secretome alterations and impaired monocyte migration in PWH and to link these findings to cellular dysfunction. Second, while AKG was used as a proxy metabolite in mechanistic assays, the broader data, including scRNAseq-defined metabolic clusters, plasma metabolomics meta-analysis, and GEM predictions, consistently converged on the Glycolysis–TCA anaplerosis axis, underscoring the robustness of our conclusions. Finally, the mechanistic assays with DM-AKG and pooled PWH serum recapitulated the high-throughput multi-omics signatures, strengthening the causal link between metabolic rewiring and macrophage dysfunction in PWH.

In summary, our integrative multi-omics and functional analyses reveal that long-term treated PWH exhibit profound immunometabolic rewiring characterized by impaired monocyte migration, altered macrophage polarization, and disrupted Glycolysis–TCA anaplerosis axis. These changes promote oxidative stress, metabolic inflexibility, and an exhausted macrophage phenotype, which may contribute to chronic immune activation and persistence of HIV reservoirs. Importantly, the convergence of high-throughput datasets with mechanistic assays underscores the central role of immunometabolic perturbations in shaping monocyte– macrophage dysfunction. Together, these findings highlight potential therapeutic opportunities to restore immune function by targeting metabolic pathways that underlie HIV-associated immune dysregulation.

## Materials and Methods

### Study population

The study population for this study includes people living with HIV (PWH) on long-term suppressive therapy (n=36) and gender and age-matched people without HIV (PWoH, n=18) from the **CO**penhagen **COMO**rbidity (COCOMO) cohort (38). The PHW initiated their antiretroviral therapy at a low CD4^+^ T-cell count median (IQR) 194 (132-300) and successful viral suppression immune reconstitution [median (IQR) CD4^+^ T-cell count 660 (550-860)]. The PWH were on treatment for a median (IQR) duration of 16 (9-18) years. The clinical and demographic data are presented in Table S1. For microenvironment MDM differential assays, pooled serum was obtained from PWH (n=6) on therapy and PWoH (n=6). Healthy blood donor samples were obtained from Karolinska University Hospital, Huddinge.

### Ethical clearances

Ethical approval was obtained by the Regional Ethics Committee of Copenhagen (COCOMO: H-15017350) and Etikprövningsmyndigheten, Sweden (Dnr: 2022-01353 and 2020-00654). Informed consent was obtained from all participants and delinked before analysis.

### Meta-analysis for untargeted metabolomics

We used our previous untageted plasma metabolomics (10, 39), from two different cohort i.e. people living with HIV (PWH) with ART (n=275), and people without HIV (PWoH) (n=100) from India, Cameroon, Sweden and Denmark. Metabolomics data were first log2-transformed and then batch-corrected to mitigate batch-specific technical variations. The batch correction was performed using the ComBat() function from the R package sva v3.42.0.(40) R package limma v3.42.2 was used for differential expression analysis.(41) P values were adjusted using Benjamini-Hochberg (BH) correction. Following the batch correction and normalization we used 536 metabolites to identify the signature of the PWH on ART.

### Single-cell RNA sequencing (scRNAseq)

Monocytes were isolated through the negative selection using EasySep™ Human Monocyte Isolation Kit (Cat#19359, STEMCELL Technologies, Canada) from PWH (n=3) and PWoH (n=1) and immediately processed for generation of Gel Bead Emulsions (GEMs) in Chromium controller, followed by library construction according to the Chromium™ Next GEM Single Cell 3′ reagent kit v3.1 (10x Genomics, USA). The libraries were run on the Illumina Nextseq 2000 P3 (Illumina, San Diego, CA).

### Data alignment and normalization

The raw fastq files were aligned and quantified using the count module from Cell Ranger v6.1.2(42). The human genome version GRCh38.96 obtained from Ensembl was used as the reference. The count matrices of PWH samples were merged using the module aggr from Cell Ranger. All the further downstream analyses of the count matrix were performed using R package Seurat v4.3.0(43). Firstly, the data quality was checked based on the total number of genes detected per cell and mitochondrial gene percentage. Cells with less than 200 genes were detected, genes expressed in less than three cells, and cells with more than 25% mitochondrial genes were filtered out. A total of 4477 cells remained after filtering. The ScaleData() function was used to scale down the count matrix, and expression variation due to cell cycle scoring was regressed. The data were then normalized using NormalizeData() function with default settings.

### Cell clustering and cluster annotation

The dimensionality of the count matrix was reduced using RunPCA() function, and corresponding principal components were computed. FindNeighbors() and FindClusters() functions were used with default settings to generate shared nearest neighbor (SNN) graph and clusters of cells by SNN modularity optimization-based clustering algorithm. Further, UMAP dimensionality reduction was performed using RunUMAP() function on the first 30 principal components, and clusters of cells were visualized. Cell types were identified by corresponding canonical cell type marker gene expression. The analysis identified a monocyte cell population (3818/4477), NK cells (544/4477), pDCs (24/4477), and cDCs (27/4477). There was also a small percentage of dying cells (27/4477) in the total cell population. The monocyte cell population was then selected and re-executed in the clustering process, as mentioned before. The monocyte-specific cell clustering identified six sub-clusters. FindAllMarkers (min.pct = 0.25, logfc.threshold = 0.25) function was used to identify marker genes specific to each cluster, and the top 15 markers of each cluster based on avg_log2FC were selected and visualized in heatmap created using DoHeatmap(). Differential expression analysis was performed using the function FindMarkers(min.pct=0.1, min.diff.pct=0.2, logfc.threshold = 0.2) and Wilcoxon Rank sum test. Pathway enrichment analysis was performed using enrichr module from the tool gseapy v0.10.5(44). The KEGG pathway gene set was used as a reference.

### Trajectory inference from single-cell data

R package Slingshot v2.7.0 was used for trajectory inference analysis from scRNA-seq data(45). The global cell lineage structure was identified using the function getLineages(). The function creates links between nearby cell clusters using a minimum spanning tree (MST) and finds trajectory through the links that denote cell lineages. Monocyte cell cluster 1 was used as starting point when calculating the path.

### Contextualizing metabolic model using MDM RNAseq and FBA

GSMM were contextualized specifically for PWH and PWoH groups using the task-driven Integrative Network Inference for Tissues (tINIT) algorithm.(46) The generic human genome-scale metabolic model obtained from the metabolic atlas was used as a reference.(47) Average normalized expression (TPM) values for each patient group were used as input. Genes with TPM < 1 were filtered out to avoid lowly expressed genes. Flux balance analysis was performed using RAVEN toolbox v2.7.11(48) and gurobi v10.0.1 solver (Frontline Systems Inc., USA).

### High throughput plasma secretome analysis by Olink® Explore

We used Olink® Explore, a multiplex immunoassay protein biomarker platform that detects approx 3000 proteins in plasma. The platform uses Proximity Extension Assay (PEA) technology and a readout based on Next Generation Sequencing (NGS) in Illumina NovaSeq 6000. Normalized protein expression (NPx) values obtained from the Olink platform were used for the analysis. Mann-Whitney U test from R function wilcox.text was used to find differentially abundant proteins. Multiple hypothesis correction using the Benjamini-Hochberg method was performed after Mann-Whitney test.

### Monocyte-derived macrophage (MDM) differentiation

PBMCs were seeded at 5×10^6^ cells/mL in RPMI 1640 (R2405, Sigma Aldrich) and incubated for two hours. Afterward, cells were cultured in full RPMI 1640 (R2405, Sigma Aldrich) media [10% FBS (10500-064, Gibco), 5% human AB serum (H3667, Sigma Aldrich), 1% penstrep (P4333, Sigma Aldrich), 10mM HEPES (H0887, Sigma)] containing 10ng/mL GM-CSF (300-03, PeproTech) for six days. Media was changed after three days, and experiments were performed on day seven for the MDM phenotype. MDM differentiated with donor or patient serum was previously heat-inactivated and substituted with the 5% human AB. When polarizing these MDM, we used 100 ng/ml of LPS from Escherichia coli (026:B6) (L5543, Sigma Aldrich) and 50 ng/ml of IFN-g (300-02, PeproTech) for M1 or 20 n/ml of IL4 (#200-04, PeproTech) for M2. For experiments involving patient serum, all commercial serum components, including FBS and human AB serum, were replaced with 5% pooled, heat-inactivated patient serum. No additional growth factors were added in this condition. In differentiation protocols involving dimethyl-α-ketoglutarate (DM-aKG), the compound was added at a final concentration of 1 mM throughout the 7-day differentiation period in conjunction with the standard MDM protocol.

### Glutamate-Glo assay

The Glutamate-Glo™ Assay (J7021, Promega) was performed according to the manufacturer’s instructions to measure intracellular glutamate levels in MDMs. Glutamate concentrations were normalized to cell number per well, as determined by crystal violet staining.

### MDM bulk transcriptomics

Following differentiation to MDM, RNA was extracted using RNeasy Mini Kit (Qiagen, Germany) and subjected to bulk RNA sequencing. The library was prepared using the SMARTer Total Stranded RNA-seq, Pico input mammalian (Takara Bio, USA). Samples were sequenced on NovaSeq6000 (NovaSeq Control Software 1.7.5/RTA v3.4.4) with a 151nt (Read1)-10nt(Index1)-10nt(Index2)-151nt(Read2) setup using ‘NovaSeqXp’ workflow in ‘S4’ mode flowcell. The Bcl to FastQ conversion was performed using bcl2fastq_v2.20.0.422 from the CASAVA software suite, followed by downstream analysis. The raw fastq files were processed using nf-core RNAseq pipeline.(49) Human reference genome GRCh38 obtained from Ensembl was used for alignment. Gene level counts data of protein-coding and lincRNA genes were considered for all the downstream analyses. Principal component analysis was performed using the R package PCAtools v2.6.0(50). R package DESeq2 v1.34.0(51) was employed for differential expression analysis. Pathway enrichment analysis was performed using the tool GSEA v4.1.0(52). The KEGG pathway gene set, which is comprised of pathways belonging to metabolism, environmental information processing, and organismal systems, was used for pathway enrichment.

### MDM single-cell type quantitative proteomics

Single-cell type quantitative LC-MS/MS-based proteomics was performed as described previously.(53) The data was first subjected to normalization using the R package NormalyzerDE v1.12.0(54). Quantile normalized data were used for further downstream analysis. Missing value imputation was done using the KNN algorithm from the R package impute v1.68.0.(55) Principal component analysis was performed using the R package PCAtools v2.6.0.(50) The batch correction was performed using the ComBat() function from the R package sva v3.42.0.(40) R package limma v3.42.2 was used for differential expression analysis.(41) P values were adjusted using Benjamini-Hochberg (BH) correction.

### Isolation of MDM by Fluorescence Associated Cell Sorting (FACS)

The MDMs were sorted using FACSAria Fusion (BD Biosciences). The cells were only sorted based on the forward and side scatter (FSC/SSC) parameters. Single cells were collected in 96-well Lo-Bind plates (Eppendorf, Hamburg, Germany) containing 5μL of 100mM triethylammonium bicarbonate (TEAB) per well. Cells from each donor were isolated into a single plate, so 1152 macrophages were collected. For a carrier proteome, 100 cells from each donor were sorted into 24 wells. All single-cell and carrier plates were centrifuged for a minute after the sorting to ensure that the cells were submerged in the buffer.

### Single-cell quantitative proteomics (SCqP) sample preparation

Proteins were extracted by carrying out freezing and thawing cycles four times. The plates were frozen by submerging them in liquid nitrogen for 2 min and immediately heated at 37°C for another 2 min. Extracted proteins were denatured by heating at 90°C for 5 min, and plates were centrifuged at 1000rpm for 2min. One µL of 25ng/µL sequencing grade trypsin (Promega, Madison, WA) in 100 mM TEAB was added per well of the single cell plates using a MANTIS^®^ automatic dispenser (Formulatrix, Bedford MA). In the carrier proteome plates, 2 µL of trypsin was added to each well. All plates were incubated at 37°C overnight.

TMTpro (16plex) reagents were used in the study, excluding channels 127N, 127C (to prevent cross-contamination from the carrier channel), and 134N. For labeling, 1 μL of the respective TMTpro reagent dissolved in dry acetonitrile (ACN) at a concentration of 10 μg/μL was dispensed in the wells using the MANTIS robot. After incubation at room temperature (RT) for 1h, the reaction was quenched by dispensing 1μL of 5% hydroxylamine (Sigma) to each well and incubating the plates at RT for 15 min. Carrier proteome was labeled with TMTpro 126 channel. From the carrier plates, six labeled wells of each donor were pooled (72 wells and 7200 cells in total) and distributed equally between all sets. Each TMT set contained one cell from each donor and a carrier proteome. In total, 36 such TMT sets were prepared. The sets were combined into sample vials with a glass insert (TPX snap ring vial from Genetec, Sweden) using a 10μL glass syringe (VWR, Japan), always starting with the carrier proteome, to minimize sample loss from single cells; and dried in vacuum (Concentrator Plus, Eppendorf). The dried TMTpro-labelled peptides were resuspended in 7 μL of 2% ACN, 0.1% formic acid (FA) before LC-MS/MS analysis.

### Reverse-phase liquid chromatography-tandem mass spectrometry (RPLC-MS/MS)

Peptides reconstituted in solvent A (2% acetonitrile and 0.1% formic acid in water) were injected onto a trap column (Acclaim PepMap 100, 2cm × 75μm, 3μm,100 Å, Thermo Fisher Scientific) and separated on a 25cm long EASY-Spray C18 column (ES802A, Thermo Fisher Scientific) installed in an Ultimate 3000 UHPLC (Thermo Fisher Scientific). The chromatographic separation was carried out at 300 nL/min, applying an organic modifier gradient from 4% to 5.5% in 29 minutes, and then further increased to 6% B in another minute. The flow rate was then reduced to 100 nL/min, and a linear gradient was applied where %B was increased from 6% to 27% in 100 min.

LC-MS/MS data were acquired on an Orbitrap Fusion Lumos Tribrid mass spectrometer (Thermo Fisher Scientific, San José CA), using nano-electrospray ionization in positive ion mode at a spray voltage of 1.9 kV. Data-dependent acquisition (DDA) mode parameters were set as follows: isolation of the most intensive precursors in full mass spectra for 2 s at 120,000 mass resolution in the *m/z* range of 350−1500, the maximum allowed injection time of 100 ms, automatic gain control (AGC) target of 400,000, and dynamic exclusion for 45s. Precursor fit of 70% in a window of 0.7 Th was applied. Precursor isolation width was set as 0.7 Th with higher-energy collision dissociation (HCD) energy 35% at a resolution of 50,000, maximum injection time of 150 ms in a single microscan, and AGC target of 50,000.

Obtained raw files were analyzed on Proteome Discoverer v2.5 (Thermo Fisher Scientific). Proteins were searched against both human (SwissProt) and HIV databases with MS Amanda 2.0, allowing for up to two missed cleavages. Mass tolerance for the precursor and fragment ions were set as 10 ppm and 0.05 Da, respectively. Oxidation of methionine, deamidation of asparagine and glutamine, and TMTpro of lysine and N-termini were set as variable modifications. The PSMs were filtered at a target false discovery rate of 1% (strict) and 5% (relaxed), with validation based on q-values. The correction was applied to account for known isotopic impurities in the TMT batch used. Normalization was done based on the total peptide amount. Normalized protein abundances were extracted from the search results.

### Targeted Metabolite Profile

The targeted metabolomics for amino acids was performed using LC-MS/MS method with reference amino acids as control at the Swedish Metabolomics Centre (Umeå, Sweden) as described by us (10). Absolute quantification of the TCA-cycle intermediates was measured by gas chromatography coupled to triple-quadrupole analyzers operating in tandem mass-spectrometer (GC-QQQ-MS) at Swedish Metabolomics Centre, Umea with an 11-point calibration curve (cis-aconitic acid, a-keto-glutaric acid, citric acid, fumaric acid, glucose, glucose-6-phosphate, isocitric acid, lactic acid, malic acid, shikimic acid, succinic acid, sucrose, urea) as described by us previously (7).

### Quantitative real-time polymerase chain reaction (qRT-PCR)

RNA was extracted from MDMs using TRI Reagent (Zymo Research) and Direct-zol™ RNA MiniPrep kit (Zymo Research) and cDNA synthesis by the SuperScript™IV reverse transcriptase (ThermoFisher Scientific). qRT-PCR was conducted using KAPA SYBR Fast universal master mix (Roche) on ABI7500F, according to the manufacturer’s protocol. Primers were synthesized from Integrated DNA Technologies, Inc. Germany, using the primer sequences listed in Table S11. Gene expression analysis was performed using the ΔΔCT method.

### Phagocytosis assay

MDMs were prepared, and on day 7, MDMs were seeded at 30000 cells/well in a sterile black well plate, letting them attach for 1h in the incubator. According to the manufacturer’s instruction, the phagocytosis assay was performed using Vybrant™ Phagocytosis Assay Kit (V-06694, Thermo Fisher Scientific).

### Monocyte migration assay

Monocytes were isolated from PBMCs (6 PWoH and 6 PWH) using the EasySep Human monocyte enrichment kit without CD16 depletion (StemCell Technologies, Canada) according to the manufacturer’s instructions. Cell migration was performed in duplicate using 24-well Transwell inserts with 5.0 µm pore polycarbonate membrane filters (Corning, USA). Monocytes (200,000 cells in 100 μl of RPMI/0.1% human serum albumin) were added to the upper chamber of the insert, and 600 μl of RPMI/0.1% human serum albumin was added to the lower chamber. CCL2 (20 ng/ml; BioLegend, USA) was also added to the lower chamber. The plates were incubated at 37°C and 5% CO_2_ for 3 hours. After that, the cells that migrated to the lower chamber were collected and counted using flow cytometry (FACSVerse™, BD Bioscience, USA). Counting beads (Invitrogen, USA) were added to each sample as a standard. Data was analyzed with FlowJo, version 10.9 (TreeStar, Inc).

### Analysis of macrophage mitochondria oxygen consumption by Seahorse XF Cell MitroStress Test

For this experiment, we used positive isolation of monocytes with EasySep™ Human CD14 Positive Selection Kit II (StemCell Technologies, Canada). Monocytes were then seeded at 150000 CD14^+^/well in 96-well Seahorse XF Cell Culture Microplate using RPMI supplemented with 2% pooled PWoH or PWH serum and 20 units/mL penicillin and 20 mg/mL streptomycin (Sigma, USA) under 5% CO_2_ and 37°C temperature. Differentiation from monocytes to macrophages took seven days. Oxygen Consumption Rate (OCR) was measured using the XFe96 Analyzer (Agilent Technologies) as described by the manufacturer. Macrophages were pre-incubated with Seahorse XF RPMI supplemented with 1mM pyruvate, 2mM glutamine, and 10mM glucose. The instrument was programmed to sequentially inject compounds into wells of the Seahorse culture plates in the following order. First, 1µM of Oligomycin for ATP production inhibition, followed by 0.5 µM of FCCP to disrupt the potential of the mitochondrial membrane, and finally 1µM of Rotenone and 1µM antimycin A for O2 consumption inhibition. Data obtained were first analyzed by Agilent Seahorse Wave Desktop software (Agilent Technologies) and normalized to cell numbers per well (quantified by crystal violet staining) and to baseline OCR levels (last measure before the first injection). OCR data is shown as the mean percentage after baseline normalization (baseline levels defined as 100%) and further normalized with the number of cells.

### Macrophage polarization phenotyping by flow cytometry

Macrophage polarization expression markers were analyzed on MDM using flow cytometry. Macrophages were stained for extracellular receptors using anti-CD1a (HI149) (748853, BD Bioscience), anti-CD163 (GHI/61) (741402, BD Bioscience), anti-CD64 (10.1) (740300, BD Bioscience), anti-CD200R (OX-108) (566344, BD Bioscience), anti-HLA-DR (L243) (335830, BD Bioscience), anti-CD3 (UCHT1) (562280, BD Bioscience), anti-CD56 (B159) (562289, BD Bioscience), anti-CD195 (2D7/CCR5) (556889, BD Bioscience),anti-CD86 (IT.2.2) (305429, Biolegend). Staining was complemented with viability dye (L34976, Invitrogen). Macrophages were permeabilized with Cytofix/Cytoperm™ Fixation/Permeabilization Kit (554714, BD Bioscience) and then stained with intracellular staining using anti-CD68 (Y1/82A) (565594, BD Bioscience). Data was acquired with BD FACS Symphony flow cytometer (BD Bioscience). Data were analyzed using FlowJo, version 10.8.1 (TreeStar, Inc) and Prism 9 (GraphPad Software, Inc).

### HIV-1 infection assay

We first differentiated monocytes from healthy donor buffy coats from Karolinska University Hospital Huddinge into macrophages. This was achieved using 20 ng/ml M-CSF (300-25, PeproTech) for seven days. The macrophages were then infected with HIV-1_ADA-M_ obtained from the NIH AIDS Reagent Program (MOI = 10). Every 48h, half of the media was replenished for 14 days. Supernatants were collected and filtered through a 0.20 µM filter on day 7 and day 14 post-infection. For the macrophage infection, cells were exposed to HIV-1_ADA-M_ (MOI = 10) for 24 hours, after which the media was changed every 24 hours. Macrophages infected with HIV-ADA were stained with viability dye (L34976, Invitrogen). Macrophages were permeabilized with Cytofix/Cytoperm™ Fixation/Permeabilization Kit (554714, BD Bioscience) and then stained with intracellular staining using HIV-1 core antigen-FITC, KC57 (6604665, Beckman Coulter). Data was acquired with BD FACS Fortessa flow cytometer (BD Bioscience). Data were analyzed using FlowJo, version 10.8.1 (TreeStar, Inc) and Prism 9 (GraphPad Software, Inc).

## Acknowledgments

The authors acknowledge support from the National Genomics Infrastructure in Genomics Production Stockholm, funded by Science for Life Laboratory, the Knut and Alice Wallenberg Foundation and the Swedish Research Council, and SNIC/Uppsala Multidisciplinary Center for Advanced Computational Science for assistance with massively parallel sequencing and access to the UPPMAX computational infrastructure. The authors would also like to acknowledge the support from Proteomics Biomedicum and the core facility for Bioinformatics and Expression Analysis Karolinska Institute. The study is funded by the The Swedish Research Council grants 2021-01756 to UN. Karolinska Instititute Consolidator Grants (2-117/2023) to UN. SDN acknowledged the support received from Novo Nordic Foundation, Lundbeck Foundation, Augustinus Foundation, Region Hovedstaden, and Rigshospitalet Research Council.

## Declaration of generative AI and AI-assisted technologies in the manuscript preparation process

During the preparation of this work the author(s) used ChatGPT v5.0 and Microsoft Copilot for language correction. After using this tools, the author(s) reviewed and edited the content as needed and take(s) full responsibility for the content of the published article.

